# No paternal effects in a sperm-dependent, naturally clonal fish

**DOI:** 10.1101/2025.07.11.664288

**Authors:** Ulrike Scherer, Sean M. Ehlman, David Bierbach, Jens Krause, Max Wolf

## Abstract

Paternal effects, i.e., effects of males on the phenotypes of their offspring that are not caused by the integration of male genetic material, are increasingly recognized as a potentially significant source of phenotypic variation across taxa - even in the absence of paternal care. Gynogenetic systems, which rely on sperm to trigger embryogenesis without incorporating male genetic material, provide a powerful way to experimentally isolate potential paternal effects from effects caused by the integration of male genetic material; up to now, however, paternal effects remain largely unexplored in these systems. Here, we test for paternal effects in the gynogenetic Amazon molly (*Poecilia formosa*): a naturally clonal, all-female species with no parental care. Using a highly controlled breeding experiment involving 59 Atlantic molly males (*P. mexicana*) and 57 Amazon molly females, we generated 169 broods and 2,966 offspring. While males were drawn from a naturally variable stock population, females – next to being genetically identical – were highly standardized for age, size, descent, and developmental experience. We asked whether male identity or body size predicted offspring size – a key offspring phenotypic trait. We also asked whether male identity or body size predicted brood size. While we found substantial variation in both offspring size and brood size, we found no evidence for paternal effects on either trait. Next to providing an experimental test for paternal effects in a gynogenetic system, our results also strengthen the Amazon molly’s status as a model species for studying – in a highly controlled fashion – the developmental emergence of phenotypic variation.

## INTRODUCTION

Parental effects – defined as any influence of parental genotype or phenotype on offspring phenotype that is not attributable to the integration of genetic material (1,2) – play an important role in shaping phenotypic variation across generations. While research on maternal effects has long dominated this field, receiving substantial empirical and theoretical attention (3–5), paternal effects have historically been overlooked (6). For decades, they were largely dismissed as relevant only in species where males provide direct care (7,8). However, a growing body of research now challenges this view, revealing that fathers can influence their offspring’s phenotype even in the complete absence of paternal care. Sperm and ejaculates can contain many non-genetic components – such as proteins, lipids, or epigenetic modifiers - that can influence embryonic development (reviewed in (1)). Moreover, females may adjust reproductive allocation in response to male traits, such as body size, through mechanisms like cryptic female choice, thereby causing an indirect link between males and offspring phenotypes (1,9,10). Thus, paternal effects are potentially more widespread and more influential than previously assumed, with recent evidence spanning taxa from insects to mammals - even in species where males never interact with their young (7).

Gynogenetic unisexual vertebrates, i.e. systems with sperm-dependent parthenogenesis, provide a powerful opportunity to experimentally isolate paternal effects from effects that are caused by the integration of male genetic material: while females in these species require sperm from a closely related sexual species to trigger embryonic development, the genetic material of the male does not contribute to the offsprings’ genotypes (11). Using the power of such a system, we here focus on the Amazon molly (*Poecilia formosa*) - a gynogenetic freshwater fish that emerged from a single hybridization event between the Atlantic molly (*P. mexicana*) and the sailfin molly (*P. latipinna*) estimated between 120,000 – 280,000 years ago (12–15). As a gynogenetic species, Amazon mollies require sperm from one of the two above mentioned parental species or other closely related poecilid species to trigger embryonic development (16). However, the male’s genetic material is not incorporated into the offspring’s genome (8–10), except in rare cases of paternal introgression (15,17,18)). We can thus experimentally isolate potential paternal effects – that is, effects that are not caused by the integration of the genetic material of the male. Up to now, the potential role of paternal effects in Amazon molly reproduction has not been assessed (but see (16) for a recent study that found an effects of male species on female reproductive success in the Amazon molly).

Here, we test for paternal effects of Atlantic molly males on offspring phenotype (offspring size at birth) in Amazon mollies. Over 34 weeks (i.e., eight months), using tightly controlled laboratory breeding, we paired 59 Atlantic molly males with 57 Amazon molly females in a fully randomized design, resulting in 169 broods and 2,966 offspring. While females were standardized for age, size, descent, and developmental experience to minimize maternal variation, males were randomly drawn from a diverse stock population, thereby preserving natural phenotypic variation among males. Critically, each male and female contributed to multiple reproductive events (approximately three broods per individual), allowing us to robustly detect potential paternal effects. We asked (i) whether male identity predicts offspring size at birth and (ii) whether male body size, often used as a proxy for male quality (19,20), predicts offspring size at birth. Beyond this, we also tested whether male identity or body size influenced brood size. While brood size itself is not an offspring trait and thus not a paternal effect in the strict sense, it remains biologically relevant: brood size and offspring size typically trade off, as larger broods often come at the cost of smaller individual offspring (21–25). Any influence of male identity on brood size could therefore represent an indirect pathway through which males affect offspring phenotypes.

## METHODS

### Study species and holding conditions

Atlantic molly males and Amazon mollies females used in our experiment originate from stock populations kept at Humboldt-Universität zu Berlin (Berlin, Germany). Both stocks are descendants from wild fish caught in Mexico. All fish were housed according to the recommendations outlined in (26), i.e., stock populations were kept in 200L tanks with approx. 50 individuals per tank under the following holding conditions: 12:12h light:dark cycle, air temperature control at approx. 24±1°C, weekly water changes. Fish were fed twice a day for 5 days a week with commercial powder food (Sera vipan baby).

Amazon mollies are live-bearing (i.e., developing offspring are carried internally), and like other poeciliid females, they follow a reproductive cycle characterized by a brief fertile period of about 2-3 days, occurring at the onset of sexual maturity and right after parturition, i.e., right after giving birth. This fertile period is followed by a roughly 30-day period during which females are not receptive to sperm. Fertilized eggs are carried internally until parturition, with gestation taking approx. 30 days (27,28). Although females can store sperm, there is an extreme fertilization bias of fresh sperm in poecilid fish (29). Specifically, (29) artificially inseminated guppy females (*P. reticulata*) with the sperm from one male, and after parturition, they were inseminated with the sperm from a different male. In the subsequent brood, the fertilization bias towards the fresh sperm was 98.6 ± 5.2% (mean ± SD).

All animal care and experimental protocols complied with local and federal laws and guidelines and were approved by the appropriate governing body in Berlin, Germany, the ‘Landesamt für Gesundheit und Soziales’ (LaGeSo G-0224/20).

### Experimental procedure

To launch our experiment, we placed *N* = 66 virgin, juveniles Amazon molly females (age = 42 days) – standardized for descent and experiential background (see ‘*Female standardisation*’ below) – into highly standardized conditions, that is, identical, individual 11L housing tanks equipped with Sera Biofibres for refuge and standardised feeding protocol, where they remained throughout the study. When females reached age = 70 days, we started the breeding phase (reproductive onset was observed to occur around this time in our previous work, (30)) and all females were allowed to reproduce until they were 260 days old.

Males (*N* = 67), randomly selected from a *P. mexicana* stock population as to reflect a natural range of phenotypic variation, were uniquely tagged with VIE elastomer markers for identification and then transferred to individual housing tanks (same as female tanks) at least 7 days before their first pairing. While females remained in their initially assigned tanks, males were rotated among them, allowing us to experimentally control for female identity and potential tank effects on reproductive characteristics at the same time.

The breeding was conducted in two phases. In phase 1, the pre-reproductive phase, every two weeks, an unfamiliar male (i.e., a male that has not already been in the female’s tank) was introduced into each female’s tank and remained there until the next male was rotated. Once a female gave birth to her first brood, the current male stayed with her for seven more days to guarantee pseudo-fertilisation. We then started phase 2, during which an unfamiliar male was added to the female’s tank whenever she gave birth to a new brood, i.e., when she was receptive to sperm. This male was left in the female’s tank for 7 days to trigger the development of her next brood. Each fish received 1/64 tsp of Sera vipan baby power food twice a day for 7 days a week, i.e., tanks with both a male and a female fish received 1/32 tsp and individual fish received 1/64 tsp food per feeding. Breeding tanks were visually separated from each other and spread over two recirculating tank systems (*N* = 51 and *N* = 50 tanks per system, respectively).

We checked all tanks for broods twice a day for 7 days a week. Once we found a brood, all offspring were taken out, counted and measured for total length (length from the tip of the snout to the end of the tail fin) (mean ± SE = 0.807 ± 0.001 mm). Female total length was also measured on the day they gave a birth. Additionally, we measured females for total length every time a male was introduced into their tank during phase 1 (see above) (16.2 ± 0.2 size measurements per female, mean ± SE = 3.078 ± 1.605 mm, *N* = 922 size measurements from 57 females). Males were measured for total length every time they were transferred to a female’s tank (10.9 ± 0.6 size measurement per male, mean ± SE = 3.491 ± 0.3996 mm, *N* = 612 size measurements from 59 males). Body sizes were measured using our custom-developed software. For each brood, the father was identified as the male who was in the female’s tank 30 days before parturition. On average, males produced 3.0 ± 1.3 broods per identity and females produced 3.1 ± 1.7 broods per identity. *N* = 6 males and *N* = 4 females did not reproduce.

### Female standardization and prior experimental experience

The data analyzed in the present study were collected using females that had previously (i.e. during the first 42 days of their life) participated in a different experiment (manuscript in preparation), we here briefly describe this experiment to clarify the origin and standardization of females. In this prior experiment, we tracked female behavior over the first six weeks of their life under highly standardised conditions (i.e., identical, individual tanks, standardised feeding protocol). For a duration of 2 weeks within this six-week window, half of the females were subjected to a predator treatment, i.e., they were presented with olfactory predator cues (*Crenicichla acutirostris*) and were chased for 2 min per day with a pike cichlid model. For the other half of the females, experimental conditions remained constant (control treatment). The prior experiment was run in *N* = 3 experimental blocks with approx. six weeks in between blocks. Females were standardized for their descent, i.e., they originated from *N* = 6 mothers, all of which were full-siblings (i.e., derived from a single brood), in the following referred to as maternal origin. All females were transferred to individual breeding tanks immediately after the prior experiment, i.e. when 42 days old.

In the present study, to control for potential carry-over effects of prior female treatment on the current study, we controlled all statistical models for potential effects of treatment (predator vs. control), experimental block and maternal origin on female reproduction (see ‘*Statistical analyses*’); we note that we found no effect of either factor (see **Supporting Information 1**).

### Exclusion of data points and missing data

For *N* = 27 broods, paternity could not be assigned with certainty as there was either no male in the tank 30 days prior to parturition (*N* = 6 broods, see above for details regarding paternity assignment) or two males were in the female’s tank 30 ± 2 days prior to parturition (during phase 1, where males were present in the female’s tank constantly; *N* = 21 broods). For *N* = 2 broods, offspring sizes are missing and for *N* = 3 broods, brood sizes are missing.

### Statistical analyses

Statistical analyses were performed in R version 4.2.1 (31) and the following packages: *arm* (32), *(33), ggplot2 (34), ggpubr* (35), *lme4* (36), and *sjPlot* (37). For all models, model assumptions were verified using residual and q-q plots. A complete summary of all models is provided in **Supporting Information 1**.

To test for an effect of male identity on the size of offspring and broods produced, we built two linear mixed-effects models (LMM) with either offspring size (*N* = 2966 offspring from *N* = 168 broods) or brood size (*N* = 169 broods) as response. Female size at parturition (predicted from individual von Bertalanffy growth curves, (38)) and tank system (i.e., females were housed in two different tank systems, see above) were included as covariates. We controlled for potential carry-over effects from female prior experimental treatment by further including female treatment (predator vs. control) and experimental block (1-3) as covariates (see ‘Female standardization and prior experimental experience’ above). As random terms, we included male identity, female identity, maternal origin, and brood identity (offspring model only). We did not control for female age, but female size at parturition as age and size were tightly correlated (linear mixed-effects model with predicted female size at parturition as response, female age as predictor, and female identity as random term, *N* = 168 broods from *N* = 57 females: **χ^*2*^**= 359.44, p < 0.001, R^2^ = 0.944). We assessed potential paternal effects from these models by calculating male repeatability (R), i.e., the proportion of the variation in offspring or brood size, respectively, that can be attributed to differences among males following (39). Female repeatability was calculated in the same manner. To test for an effect of male size on reproductive characteristics, we added male total length (averaged over all individual size measurements) as a predictor to the above-described models.

## RESULTS

Male identity did not explain the observed variation in the size (repeatability with 95% confidence interval = 0.000 [0.000, 0.000]) or number of offspring (repeatability with 95% confidence interval = 0.000 [0.000, 0.000]) produced (**Figure 1**). Similarly, we found no effect of male size on offspring size (*N* = 2966 offspring from 59 males, **χ^*2*^**= 1.136, p = 0.286) or brood size (*N* = 168 broods from 59 males, **χ^*2*^**= 0.066, p = 0.798) (**Figure 2**). As our study was conducted over a period of ten months (with the typical lifespan of poecilid fish in the lab being ∼2 years), and as males of unknown ages were sampled from the stock population, some males died during the experiment (*N* = 10 males). However, our results are robust with respect to the removal of offspring or broods, which were sired by males who died within two months post pseudo-fertilization (**Supporting Information 2**).

**Figure 1.**
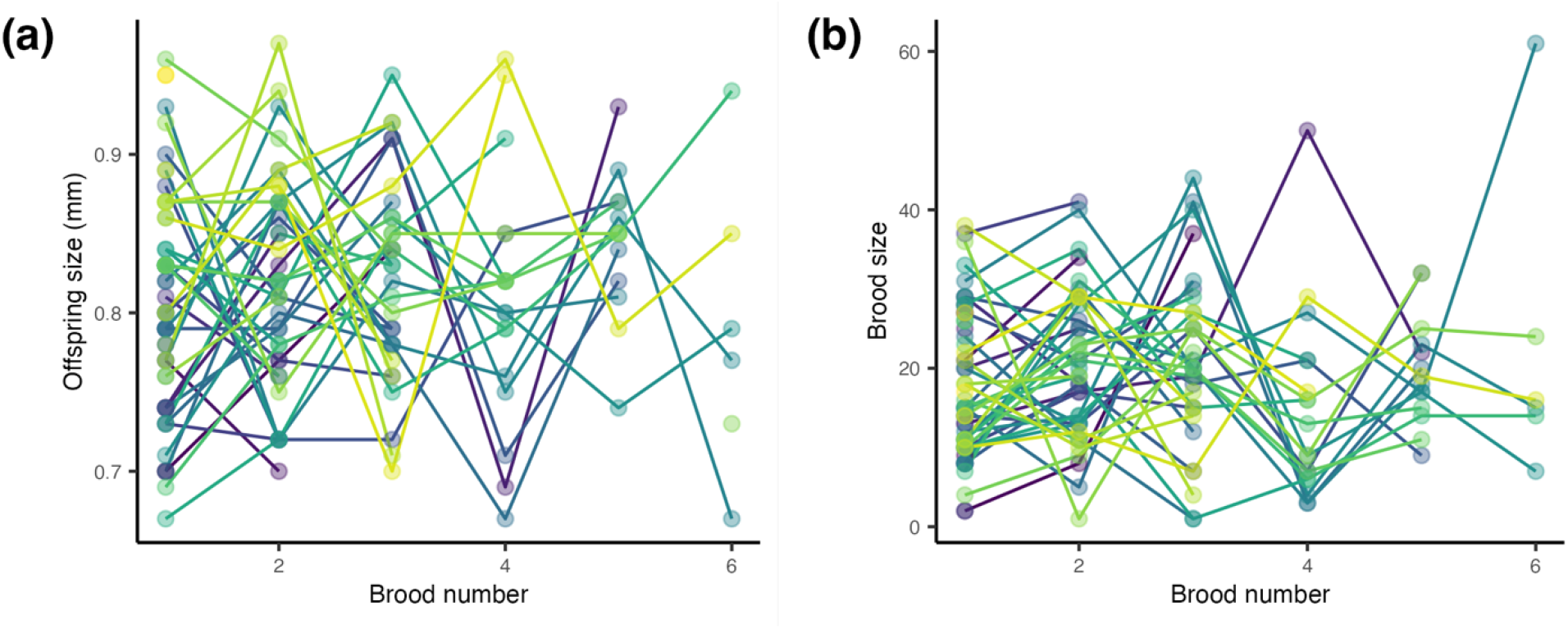
Male identity did not explain the observed variation in (a) offspring size and (b) brood size. Shown are raw data with one data point per brood, colored by male identity. Males produced at least one, but up to six broods (mean ± SD = 3.0 ± 1.3 broods per male), broods are numbered chronologically per male.

**Figure 2.**
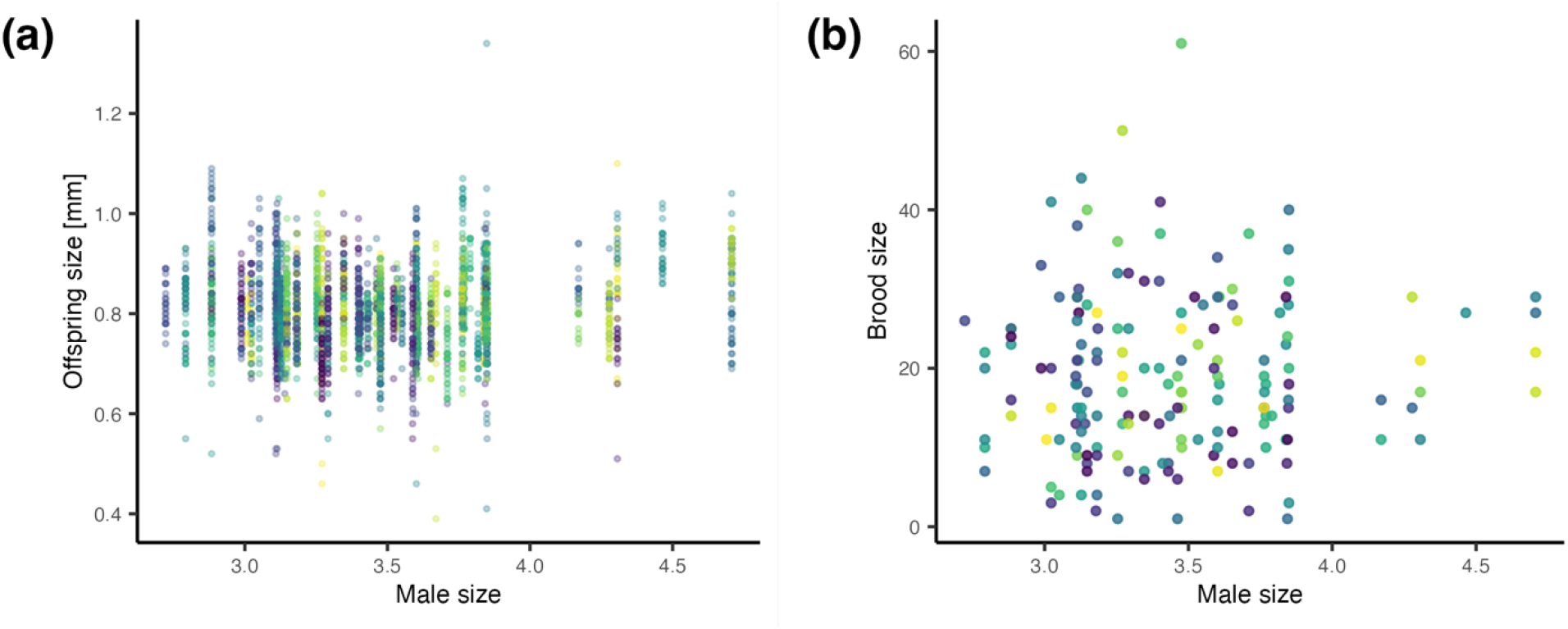
Male size did not predict (a) offspring size (*N* = 2966 offspring) and (b) brood size. (*N* = 169 broods). Shown are raw data, i.e., one data point per offspring (a) or brood (b), colored by male identity.

We note that, despite the lack of genetic differences among females and the minimization of environmental differences among them, we found significant variation in the size of offspring (repeatability with 95% confidence interval = 0.188 [0.136, 0.249]) and broods (repeatability with 95% confidence interval = 0.066 [0.043, 0.090]) produced among females. Maternal origin also explained a small, yet significant proportion of among-female variation in offspring size (repeatability with 95% confidence interval = 0.020 [0.004, 0.055]), but not brood size (repeatability with 95% confidence interval = 0.000 [0.000, 0.000]).

## DISCUSSION

We used a gynogenetic system, where males contribute neither genes nor care to reproduction, to experimentally test for potential paternal effects. Leveraging a highly controlled experimental setup and a robust repeated-measures design, we found no evidence that either male identity or male body size influenced offspring size in the Amazon molly. We also found no evidence that either male identity or male body size affected brood size. These findings were despite substantial observed variation in both offspring size and brood size, with variation in both traits being linked to female identity. While offspring size at birth is a key phenotypic trait, our findings do not rule out potential paternal effects on unmeasured aspects of offspring development (e.g., offspring behaviour, offspring growth). It will thus be interesting to investigate a broader set of offspring phenotypic traits in future studies.

Our study provides a test of a long-standing assumption in the field of reproductive biology: in gynogenetic systems, like the Amazon molly, where males contribute neither genes nor care, males are irrelevant to offspring phenotypes. By leveraging a highly controlled experimental setup with a large sample size (i.e., 59 ‘fathers’, 57 mothers, resulting in 169 broods and 2,966 offspring), we show that this assumption may hold true - at least with respect to offspring size at birth and brood size. Despite substantial natural variation among Atlantic molly males and a well-powered design capable of detecting subtle effects, neither male identity nor male body size explained variation in the size or number of offspring produced. In contrast, consistent with prior findings (30), maternal identity did significantly predict both traits, even among genetically identical, size- and age-controlled females raised under highly standardized conditions.

Our findings also carry important implications for the growing use of Amazon mollies as a model system in behavioural and developmental research. As researchers increasingly turn to clonal organisms to isolate the effects of subtle environmental differences and developmental stochasticity (30,40–48), it becomes imperative to assess potential hidden sources of variation in these species. Our study directly addresses one such source: the potential confounding role of males in a gynogenetic system. In doing so, we provide evidence that paternal effects on key reproductive characteristics (offspring and brood size) are negligible in one important study system, the Amazon molly. This finding thus strengthens the case for Amazon mollies as a powerful tool to investigate the developmental origins of phenotypic variation, including the role of random developmental noise, minute early-life experiences, and feedbacks (30,40,42,49).

In conclusion, our findings reinforce a central tenet of gynogenetic reproduction: that males in these systems serve only to trigger, but not shape, embryonic development. By ruling out paternal effects on two fundamental reproductive traits in the Amazon molly, we help clarify the boundaries of male influence in sperm-dependent parthenogenesis. At the same time, our results reinforce the Amazon molly’s status as an important clonal model system for dissecting the developmental origins of phenotypic variation - highlighting the Amazon molly not only as an evolutionary curiosity, but as a powerful experimental model for the study of individual differences.

## ACKNOWLEDGEMENTS

We gratefully acknowledge funding from the Deutsche Forschungsgemeinschaft (DFG) under Germany’s Excellence Strategy EXC 2002/1, ‘Science of Intelligence’ (project number #390523135) and DFG ‘Eigene Stelle’ grant to SME (#536703956). Members of the lab at the Humboldt University of Berlin and the Leibniz Institute of Freshwater Ecology and Inland Fisheries provided invaluable help with animal husbandry, experimental routine, data acquisition and processing; particular thanks are extended to Christopher Schutz and Ronja Leipoldt.

